# Cofilin1 oxidation links oxidative distress to mitochondrial demise and neuronal cell death

**DOI:** 10.1101/2020.09.09.289710

**Authors:** Lena Hoffmann, Marcel S. Waclawczyk, Eva-Maria Hanschmann, Manuela Gellert, Marco B. Rust, Carsten Culmsee

## Abstract

Many cell death pathways, including apoptosis, regulated necrosis and ferroptosis are relevant for neuronal cell death and share common mechanisms such as the formation of reactive oxygen species (ROS). However, which molecular signaling pathways contribute to related pathologies and how they are interconnected remains elusive.

Here, we present the role of cofilin1 in regulating mitochondrial functions and neuronal impairment. Cofilin1 deletion in neuronal HT22 cells exerted increased mitochondrial resilience, assessed by quantification of mitochondrial ROS production, mitochondrial membrane potential and ATP levels. HT22 cells deficient for cofilin1 exhibited a profound glycolytic shift to meet their energy demand in conditions of erastin and glutamate toxicity, whereas control cells were metabolically impaired and underwent ferroptosis and oxytosis, respectively. Further, cofilin1 was confirmed as a key player in glutamate-mediated excitotoxicity in primary cortical neurons isolated from cofilin1^flx/flx, CaMKIIα-Cre^ knock-out mice. Mitochondrial respiration and cell viability were significantly preserved in cofilin1^-/-^ primary neurons under conditions of excitotoxicity.

Using isolated mitochondria and recombinant cofilin1, we provide a further link to toxicity-related mitochondrial impairment mediated by oxidized cofilin1. Wildtype cofilin1 directly affected the mitochondrial membrane potential, mitochondrial ROS accumulation and mitochondrial respiration. The detrimental impact of cofilin1 on mitochondria depends on oxidation of cysteine residues at positions 139 and 147.

Our findings show that the actin-regulating protein cofilin1 acts as a redox sensor in oxidative cell death pathways of ferroptosis and oxytosis, and also promotes glutamate excitotoxicity. Oxidized cofilin1 links ROS accumulation to mitochondrial demise and neuronal cell death. Protective effects by cofilin1 inhibition are particularly attributed to preserved mitochondrial integrity and function. Thus, interfering with the oxidation and pathological activation of cofilin1 may offer an effective therapeutic strategy in neurodegenerative diseases.

## Introduction

Oxidative stress has been linked to many disorders including neurological pathologies. Recently, the term has been redefined and divided into oxidative eu- and distress, acknowledging redox regulation of specific targets in physiological signal transduction [1, 2]. Oxidative distress can induce neuronal cell death and is widely considered as a pivotal cause of cell death in neurodegenerative disorders, such as Alzheimer’s (AD) or Parkinson’s disease (PD) [3, 4]. In the last decades, pathophysiological mechanisms contributing to neuronal cell damage through different forms of regulated cell death (RCD) were studied intensively [5]. It is widely accepted that major steps of the cell death cascade comprise detrimental accumulation of intracellular calcium and formation of reactive oxygen species (ROS) [3, 6]. Moreover, different RCD paradigms converge at the level of mitochondria [7]. Mitochondria are dynamic organelles regulating the energy metabolism, calcium homeostasis and the cellular redox balance [8]. Thus, mitochondrial demise, including mitochondrial calcium overload, loss of the mitochondrial membrane potential, accumulation of reactive oxygen species and release of apoptosis inducing factor (AIF) are considered as the ‘point of no return’ upon cell death induction [9, 10]. A broad understanding of the molecular mechanism involved in transducing detrimental cell death signals to mitochondria are of great importance for future clinical implications.

In the current study, regulated cell death was induced by glutamate or erastin treatment leading to cell death mechanisms called oxytosis or ferroptosis, respectively. Oxytosis is a well-established form of regulated cell death occurring during neuronal development, as well as under pathological conditions in neurodegenerative diseases [11]. In addition, ferroptosis was defined more recently as an iron-dependent form of oxidative cell death, which can be achieved, e.g., by erastin treatment in neuronal HT22 cells [12, 13]. Both forms of cell death share very similar RCD mechanisms in neuronal cells, and both, glutamate- or erastin-induced inhibition of the cystine-glutamate (X_c_^-^)-antiporter leads to a reduction in glutathione levels by depletion of intracellular cysteine. This results in impaired activity of the glutathione peroxidase-4 (Gpx4) and activation of 12/15-lipoxygenase (LOX) and accumulation of ROS [14, 15]. In turn, dynamin-related protein 1 (DRP1) and the pro-apoptotic protein BID attain activity to translocate to mitochondria inducing mitochondrial ROS production and loss of the mitochondrial membrane potential by mitochondrial outer membrane permeabilization (MOMP) [13, 16, 17]. Finally, cytochrome c and apoptosis inducing factor (AIF) are released from mitochondria and translocate to the nucleus, where AIF is involved in the degradation of deoxyribonucleic acid (DNA) [14, 15, 18].

Cofilin1 is a member of the ADF/cofilin family of actin-depolymerizing proteins and the major representative of this family in neurons [19]. Upon dephosphorylation of serine residue at position 3 (Ser3) of the protein, it can bind to filamentous actin (F-actin) and initiate its depolymerization [20]. Moreover, it can bind to globular actin monomers (G-actin) and inhibit the nucleotide exchange from ADP-actin to ATP-actin, which is required for F-actin assembly [21]. Thus, cofilin1 can exert indirect effects on molecular mechanisms by operating on actin dynamics and it can also act as a direct participator in the apoptotic cell death cascade by recruitment of cofilin1 from the cytosol to mitochondria [22]. Importantly, cofilin1’s effects on mitochondria can be versatile, as it was shown to be involved in transducing apoptotic signaling to mitochondria upon oxidation [22, 23]. Oxidation of cofilin1 is an important posttranslational modification for the regulation of cytoskeletal dynamics (reviewed in [24]). Moreover, acting on mitochondrial dynamics via DRP1 activation was demonstrated in a cofilin1 loss-of-function approach in mouse fibroblasts [25].

How cofilin1 can also participate in non-apoptotic cell death paradigms remains to be elucidated. Therefore, we investigated the effect of cofilin1-depletion in non-apoptotic cell death induced by glutamate or erastin in neuronal HT22 cells and primary cortical neurons. The results obtained in this study demonstrate a role for cofilin1 as a redox sensor and regulator of oxytosis or ferroptosis, upstream of mitochondria, in neuronal cells.

## Materials and methods

### Cell culture

HT22 cells originate from immortalized primary mouse hippocampal neurons. HT22 cells were incubated at 37 °C and 5 % CO_2_ in Dulbecco’s modified Eagle’s high glucose medium (DMEM; Sigma-Aldrich, Munich, Germany) supplemented with 10% fetal bovine serum, 20 mM HEPES, 100 units/mL penicillin, 100 μg/mL streptomycin, and 2 mM glutamine.

For efficient cofilin 1 knockdown, HT22 cells were transfected with 15 nM cofilin1 siRNA for 48 h using Lipofectamine RNAiMAX (Thermo Fisher Scientific, Darmstadt, Germany) according to the manufacturer. Control cell were transfected with unspecific scrambled control siRNA. Following siRNA sequences were obtained from Dharmacon: scrambled siRNA (scrsiRNA, 5’-UAAUGUAUUGGAACGCAUA-3’), *Cofilin1* siRNA1 (CflsiRNA1, 5’-AGACAAGGACUGCCGCUAU-3’) and *Cofilin1* siRNA2 (CflsiRNA2, 5’-GGAAUCAAGCAUGAAUUAC-3’).

### Primary cortical neurons

Primary cortical neurons were prepared from embryonic mouse brains (E18) as described previously [18]. Dissociated neurons were seeded at a density of 45,000 cells per well onto polyethyleneimine (PEI) coated 96-well plates or 6-well plates with 550,000 cells for western blot analysis. Neuronal cultures were grown in neurobasal medium (ThermoFisher Scientific, Darmstadt, Germany) supplemented with 1.2 mM glutamine, 2 % B27 Plus supplement (ThermoFisher Scientific, Darmstadt, Germany), 100 U/mL penicillin and 100 μg/mL streptomycin. Every three to four days, half of the medium was replaced by fresh supplemented neurobasal medium. Glutamate treatment (25 μM) was conducted at day 30 in culture (DIV 30) for 24 h. NMDA-antagonist MK801 (Merck KGaA, Germany) was added as a control at a concentration of 10 μM simultaneously to glutamate addition. Rho activator II CN03 (Cytoskeleton, Denver, USA) was applied at a concentration of 1 μg/mL 3 h prior to glutamate treatment.

### Cofilin1^flx/flx^ mice

Genetically modified mice expressing Cofilin1 allele with exon 2 flanked by loxP sites were used as controls (Ctrl) [26]. Cofilin1 knockout was achieved by expression of the Cre enzyme capable of recognizing loxP sites and thus specifically deleting exon 2 of the cofilin1 gene region, resulting in a non-functional gene product. Since a systemic knockout of cofilin1 is embryonically lethal [27], Cre expression is under the control of a CaMKIIα-promotor to specifically delete cofilin1 in excitatory neurons for the forebrain including cerebral cortex neurons [28].

### RT-PCR

HT22 cells were plated in a 6-well plate (170,000 cells/well). Total RNA was isolated by using InviTrap Spin Universal RNA Mini Kit (Stratec Biomedical, Birkenfeld, Germany) 48 h after siRNA transfection. SuperScript III One-Step RT-PCR System (Thermo Fisher Scientific, Darmstadt, Germany) was used to perform reverse transcription PCR (RT-PCR) and specific oligonucleotides (*Gapdh* (399 bp) forward 5’-CGTCTTCACCACCATGGAGAAGGC-3’ and reverse 5’-AAGGCCATGCCAGTGAGCTTCCC-3’ and *Cofilin1* (146 bp) forward 5’-GCCAACTTCTAACCACAATAG-3’ and reverse 5’-CCTTACTGGTCCTGCTTCC-3’). The amplification products were visualized by agarose gel electrophoresis after staining with ethidium bromide by illumination with UV light.

### Protein analysis

Protein extraction and Western blot analysis were performed as previously described [13]. Briefly, cells were ruptured 48 h after transfection in a protein lysis buffer containing 0.25 M D-mannitol, 0.05 M Tris base, 1 mM EDTA, 1 mM EGTA, 1 mM DTT, and 1% Triton X-100 supplemented with protease and phosphatase inhibitor cocktail tablets (Roche Diagnostics, Mannheim, Germany). Liquid N_2_ was used to devastate cell membranes to collect whole protein lysate. Insoluble fragments were removed by centrifugation at 10,000 x g for 15 min at 4 °C. The total protein amount was determined using the Pierce BCA Protein Assay Kit (Thermo Fisher Scientific, Darmstadt, Germany). Gel electrophoresis was conducted to separate proteins due to their molecular weight. Afterwards, the proteins were transferred onto a PVDF membrane (Roche Diagnostics, Mannheim, Germany). Cofilin1 and phospho-Cofilin1 (Ser3) (1:1000; Cell Signaling Technology, Danvers, USA), as well as α-tubulin as a loading control were analyzed using indicated primary antibodies (α-Tubulin antibody 1:10,000; Sigma-Aldrich, Munich, Germany). After incubation with the appropriate secondary HRP-labeled antibody (Vector Laboratories, Burlingame, CA, USA) Western Blot signals were detected by chemiluminescence with Chemidoc system (Bio-Rad, Munich, Germany).

### Cell viability

#### MTT assay

Cell viability was assessed by a colorimetric assay based on the yellow colored MTT reagent (3-(4,5-dimethyl-2-thiazolyl)-2,5-diphenyl-2H-tetrazolium bromide, 0.5 mg/mL for HT22 cells and 1 mg/mL for primary cortical neurons; Sigma-Aldrich, Munich, Germany) which is reduced to a purple colored formazan product within a 1 h incubation period at 37 °C. The colored product can be quantified by absorbance measurement at 570 nm with a reference filter at 630 nm by FluoStar OPTIMA reader (BMG Labtech, Ortenberg, Germany).

#### xCELLigence measurement

Cell proliferation and detachment was monitored using the xCELLigence Real-Time Cell Analysis (RTCA; Roche Diagnostics, Mannheim, Germany) system as previously described [29]. Changes in the impedance are depicted as normalized cell index. HT22 cells were seeded in 96-well plates at a density of 6,000-8,000 cells per well.

### Flow cytometry

Different cellular and mitochondrial parameters of the glutamate- or erastin-induced cell death pathways were analyzed using the Guava easyCyte 6–2L flow cytometer (Merck Millipore, Darmstadt, Germany) upon harvesting adherent HT22 cells and following addition of different fluorescent dyes.

#### Cell death

Apoptotic and late necrotic cells were identified using the Annexin V-FITC Detection Kit (Promokine, Heidelberg, Germany). AnnexinV and propidium iodide (PI) staining were performed for 5 min in the dark at room temperature after harvesting the cells with trypsin.

#### Lipid peroxidation

By staining the cells with BODIPY 581/591 C11 (4,4-difluoro-5-(4-phenyl-1,3-butadienyl)-4-bora-3a,4a-diaza-s-indacene-3-undecanoic acid; Thermo Fisher Scientific, Darmstadt, Germany) oxidized lipids and membranes were detected. Following 8 h of glutamate treatment, the cells were stained with 2 μM BODIPY dye for 1 h at 37 °C.

#### Cellular reactive oxygen species formation

The cell permeable dye 2’,7’-dichlorodihydrofluorescein diacetate (H_2_DCF-DA) was used to evaluate the accumulation of cellular reactive oxygen species upon 30 min incubation of 20 μM DCF dye in DMEM without serum.

#### Mitochondrial superoxide formation

For evaluation of mitochondrial reactive oxygen species accumulation, MitoSOX Red indicator (Thermo Fisher Scientific, Darmstadt, Germany) was applied at a concentration of 1.25 μM for 30 min at 37 °C. For the MitoSOX measurement with isolated mitochondria, 10 μM of the complex III-inhibitor antimycin A (AA) was used as a positive control and incubated simultaneously with MitoSOX Red indicator at a concentration of 1.25 μM and afterwards measured with Guava easyCyte 6–2L flow cytometer (Merck Millipore, Darmstadt, Germany)

#### Mitochondrial membrane potential

Mitochondrial membrane potential was measured upon staining the cells with MitoPT TMRE Kit (ImmunoChemistry Technologies, Hamburg, Germany). Therefore, cells were incubated with 0.2 μM TMRE (tetramethylrhodamine ethyl ester) for 30 min at 37 °C. For investigation of the mitochondrial membrane potential of isolated mitochondria from adult mouse brain tissue, 40 μg mitochondria were diluted in 150 μL 1x mitochondrial assay solution (MAS: 70 mM sucrose, 220 mM mannitol, 10 mM KH_2_Cl_2_, 5 mM MgCl_2_, 2 mM HEPES, 1 mM EGTA, 0.20 % BSA, pH7.2) supplemented with 2 μM rotenone and 10 mM succinate. 50 μM of the uncoupler CCCP was used as a positive control. Mitochondria were incubated with 0.2 μM TMRE for 15 minutes and then measured with Guava easyCyte 6–2L flow cytometer (Merck Millipore, Darmstadt, Germany)

#### Mitochondrial calcium measurement

Staining the cells with the mitochondrial selective dye Rhod-2 AM (rhodamine-2 acetoxymethyl ester; Thermo Fisher Scientific, Darmstadt, Germany), allows for specific evaluation of mitochondrial calcium accumulation. Therefore, Rhod-2 AM was reduced to Dihydrorhod-2 AM and incubated at a concentration of 2 μM in DMEM without serum for 1 h.

### ATP measurement

Cellular ATP levels were measured using the ViaLight Plus Kit (Lonza, Verviers, Belgium) according to the manufacturer’s protocol. Briefly, cells were lysed, transferred to a white-walled 96-well plate and the ATP monitoring reagent was added to the cell lysate. Afterwards, the luminescence was detected with a FLUOstar OPTIMA reader (BMG Labtech, Ortenberg, Germany).

### Measurement of mitochondrial oxygen consumption rate (OCR) and extracellular acidification rate

Determination of the mitochondrial oxygen consumption rate as an indicator of mitochondrial respiration was performed using the Seahorse XFe96 Analyzer (Agilent Technologies, Waldbronn, Germany). Cells were plated in XFe96-well microplates (6000 cells/well, Seahorse Bioscience) and 1 h prior to the measurement, growth medium was replaced by the seahorse assay medium (4.5 g/l glucose, 2 mM glutamine, 1 mM pyruvate, pH 7.35). After recording three baseline measurements, four compounds were added by injection.

The compounds and final concentrations used are as follows: oligomycin 3 μM, FCCP 0.5 μM, rotenone 0.1 μM, and antimycin A 1 μM. For measuring mitochondrial respiration, 5-12 μg of isolated mitochondrial protein was measured in 1xMAS containing the complex II substrate succinate (10 mM) and the complex I inhibitor rotenone (2 μM) to focus on mainly complex II- and complex III-driven respiration. For slight attachment of the plated mitochondria at the bottom of the cell plate, a centrifugation step of the whole plate at 2,000 x g for 20 min at 4 °C was indispensable (Heraeus Megafuge 40R; Thermo Fisher Scientific, Darmstadt, Germany). Indicated compounds for the injections were used in following final concentrations: 4 mM ADP, 2.5 μg/mL oligomycin, 4 μM FCCP and 4 μM antimycin A (AA). Three basal and three measurements after each injection were recorded with the Seahorse XFe96 Analyzer (Agilent Technologies, Waldbronn, Germany)

### Mitochondrial isolation

Mitochondrial isolation of freshly dissected cortical or hippocampal brain tissue (~50 mg) was performed as previously described [18]. The tissue was embedded in 2 mL mitochondrial isolation buffer (composed of 300 mM sucrose, 5 mM TES, 200 μM EGTA, pH 7.2) and roughly homogenized with a 20G Neoject needle (Dispomed, Gelnhausen, Germany) and then sieved through a 100 μm nylon cell strainer (Corning Incorporated, Corning, NY, USA). To homogenize the tissue efficiently and extract mitochondria from the cell structure thoroughly, a cell homogenizer (Isobiotec, Heidelberg, Germany) with 1 mL gas-tight syringes (Supelco, Munich, Germany) was used to ensure a constant rate of 700 μL/min. The cell homogenizer contained a spherical tungsten carbide ball with a clearance of 10 μm to decompose the tissue but simultaneously maintain the integrity of mitochondria. The cell homogenate was transferred into 1.5 mL tubes and centrifuged at 800 x g for 10 min at 4 °C to remove cell debris. Afterwards, the supernatant was transferred into a fresh tube and centrifuged at 10 000 x g again for 10 min at 4°C (Heraeus™ Fresco™ 17 Mikrozentrifuge; Thermo Fisher Scientific, Darmstadt, Germany). The resulting pellet consists of the crude mitochondrial fraction, which was finally resuspended in MSHE-BSA (composed of 70 mM sucrose, 210 mM mannitol, 5 mM HEPES, 1 mM EGTA, 0.5 % (w/v) BSA, pH 7.2) buffer. The seahorse measurement was performed in mitochondrial assay solution, composed of 70 mM sucrose, 220 mM mannitol, 10 mM KH_2_PO_4_, 5 mM MgCl_2_, 2 mM HEPES, 1 mM EGTA, 0.2 % (w/v) BSA, pH 7.2). All steps were performed on ice or at 4 °C. Pierce™ BCA Kit was used to determine the protein amount of the mitochondrial fraction.

### Protein expression and purification

#### Cloning and mutagenesis

Human cofilin1 was amplified by PCR using specific oligonucleotides (forward: 5’-CATATGGCCTCCGGTGTG-3’, reverse: 5’-GGATCCTCACAAAGGCTTGCCCTC-3’) and cloned into the pET-15b vector (Novagen, Millipore, UK). By using site-directed mutagenesis, the cysteine residues of cofilin1 were mutated into serine residues using complementary oligonucleotides harboring the nucleotide exchanges (Cys39Ser: forward: 5’-GTGCTCTTCTCCCTGAGTG-3’, reverse: 5’-CACTCAGGGAGAAGAGCAC-3’; Cys80Ser: forward: 5’-CATAAGGACTCCCGCTATGC-3’, reverse: 5’-GCATAGCGGGAGTCCTTATC-3’; Cys139Ser: forward: 5’-CAAGCAAACTCCTACGAGGAG-3’, reverse: 5’-CTCCTCGTAGGAGTTTGCTTG-3’; Cys147Ser: forward: 5’-GACCGCTCCACCCTGG-3’, reverse: 5’-CCAGGGTGGAGCGGTC-3’) and the KOD Hot Start Mastermix (Merck, Darmstadt, Germany). The plasmids were confirmed by sequencing (Seqlab, Göttingen, Germany).

#### Expression, purification and thermal stability

The human cofilin1 WT, the mutant lacking two Cys residues (Cys139/147Ser, 2Cys → Ser) and the mutant lacking all 4 Cys residues (4Cys → Ser) were expressed as His-Tag fusion proteins in *E. coli* as described before [30]. The proteins were purified by immobilized metal affinity chromatography using the His Trap Kit from GE Healthcare Life Science, USA. Expression and purification efficiency were analyzed by SDS-PAGE using precast gels from BioRad, USA, and Coomassie staining. Proteins were re-buffered into PBS using Zeba Spin columns and the thermal stability of the proteins was analyzed by recording the emission at 600 nm over time with increasing temperature from 20 °C to 70 °C (2 °C per 3 min) using the Shimadzu UV1800.

Recombinant cofilin1 was either used in the native way, oxidized by 100 μM H_2_O_2_ incubation for 30 min, or reduced with 10 mM freshly dissolved dithiothreitol (DTT) for 30 min. The remaining elution buffer from the protein purification process was substituted by PBS using sephadex-based PD MidiTrap G-25 columns (GE Healthcare, Chicago, USA). Afterwards, protein amount was determined by a NanoPhotometer™ (Implen, Munich, Germany). The experiments were performed using 0.13-0.25 μg recombinant protein per μg mitochondrial protein and incubated for 30 - 60 minutes at room temperature and another 10 minutes at 37 °C and afterwards measured as indicated at the respective method.

### Results

#### Cofilin1 downregulation attenuates glutamate- and erastin-induced cell death

The phosphorylation and oxidation state of cofilin1 determines not only its binding capacity to F-actin [31–33], it is also essential for translocation to mitochondria [23]. In particular, dephosphorylated cofilin1 attains activity for translocation from the cytosol to mitochondria [34]. Here, mmortalized mouse hippocampal HT22 cells were used to examine the relevance of cofilin1 involvement in neuronal cell death mechanisms.

Cofilin1 involvement in cell death pathways, such as glutamate-induced oxytosis or erastin-induced ferroptosis in neuronal HT22 cells and its dependency on the cofilin1 phosphorylation status was demonstrated by Western blot analysis. Cofilin1 was dephosphorylated between 8 and 14 h after the administration of glutamate or erastin, respectively (Fig. 1 A, B). Cofilin1 protein levels started to diminish after 16 h of damaging the cells with erastin or glutamate (Fig. 1A, B). In cell death paradigms with oxidative distress as a prerequisite, a loss-of-function approach was valuable to specify the importance of cofilin1. Therefore, cofilin1 downregulation was achieved by incubation with two different siRNA sequences (si01, si02) for 48 h. The knockdown efficacy at mRNA level was detected by RT-PCR with specific primers leading to an amplification product with a size of 146 base pairs (Fig. 2 B). *Gapdh* primers were used as a loading control. At the protein level, Western blot analysis revealed an efficient knockdown of cofilin1 (Fig 2 A). At the molecular level, knockdown of cofilin1 was sufficient to prevent cellular damage after 16 hours of erastin or glutamate exposure. Particularly, the metabolic activity, representing the viability and survival of cells, revealed a protective outcome of cofilin1-knockdown upon erastin or glutamate treatment (Fig. 2 C), as assessed by MTT assay [35]. Further, deletion of cofilin1 reduced the number of AnnexinV and propidium iodide (PI) positive cells upon erastin or glutamate exposure, measured by FACS analysis, thereby confirming the protective effects of cofilin1 depletion (Fig. 2 D). For real time monitoring of the observed beneficial effects of cofilin1 silencing in living cells, the xCELLigence real-time cell analysis (RTCA) system was applied. For this approach, HT22 cells were transfected with the respective siRNA sequence, and 40 h later, cells were challenged with glutamate or erastin. Within 5 to 10 h of treatment, untransfected control cells and cells transfected with the unspecific siRNA detached from the cell plate, represented by a decline in the cell index. In line with previous findings, cell detachment of cofilin1-depleted cells was significantly attenuated (Fig. 2 E, F). These effects demonstrate a relevant involvement of cofilin1 in oxidative cell death pathways, namely oxytosis and ferroptosis.

**Figure 1.**
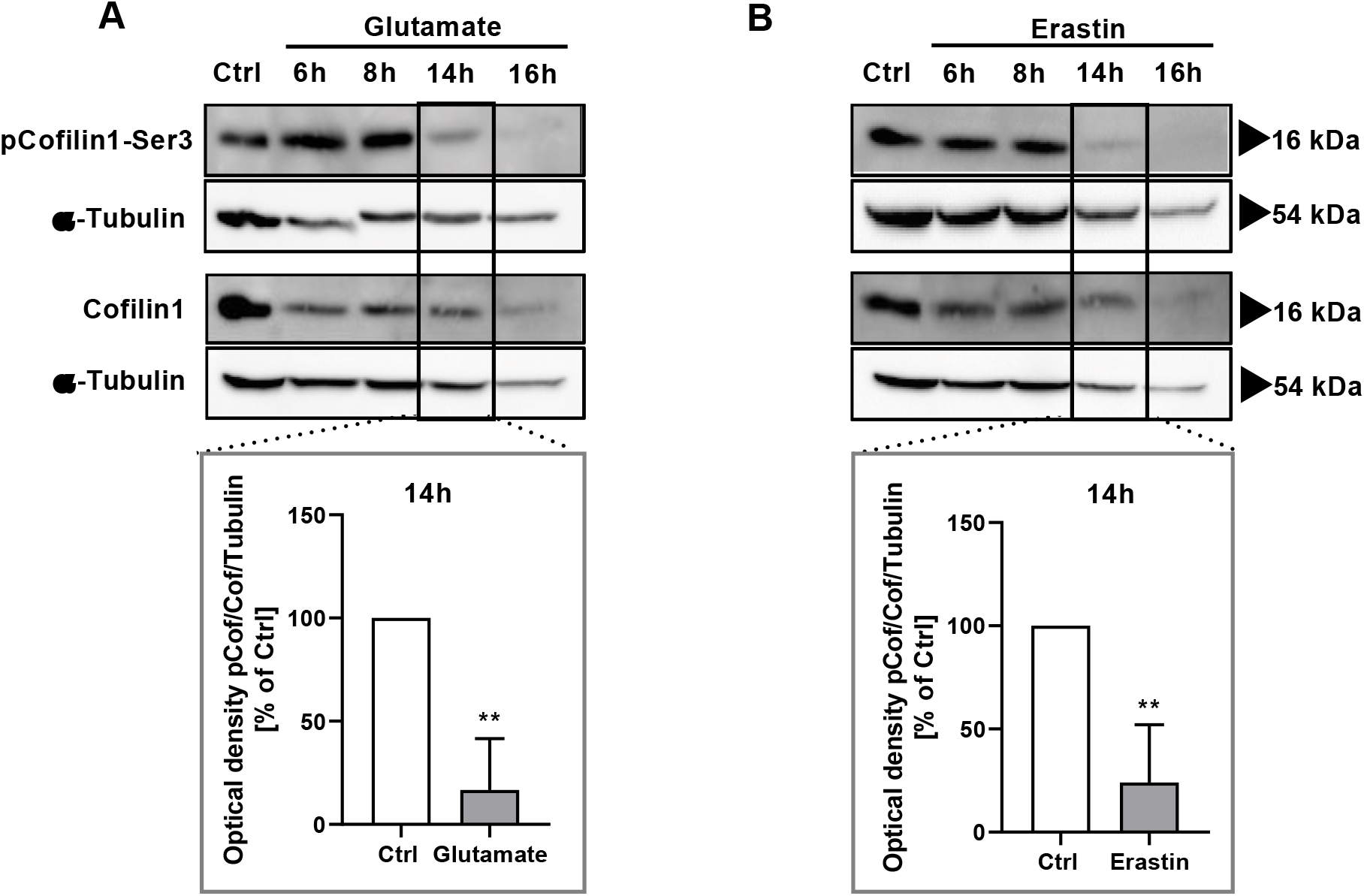
Cofilin1 is activated by dephosphorylation under conditions of oxidative stress. **A** HT22 cells were challenged with 10 mM glutamate for the indicated time and total as well as phosphorylated cofilin1 (Ser3) was analyzed via Western blot. Three blots were quantified (mean + SD). **B** Accordingly, HT22 cells were treated with 1 μM erastin for the specified time and total as well as phosphorylated cofilin1 (Ser3) was analyzed via Western blot and afterwards quantified from three independent blots. (mean + SD). Ctrl (control), pCof (phosphorylated cofilin1-ser3), Cof (cofilin1). ** p<0.01 compared to Ctrl (unpaired t-test).

**Figure 2.**
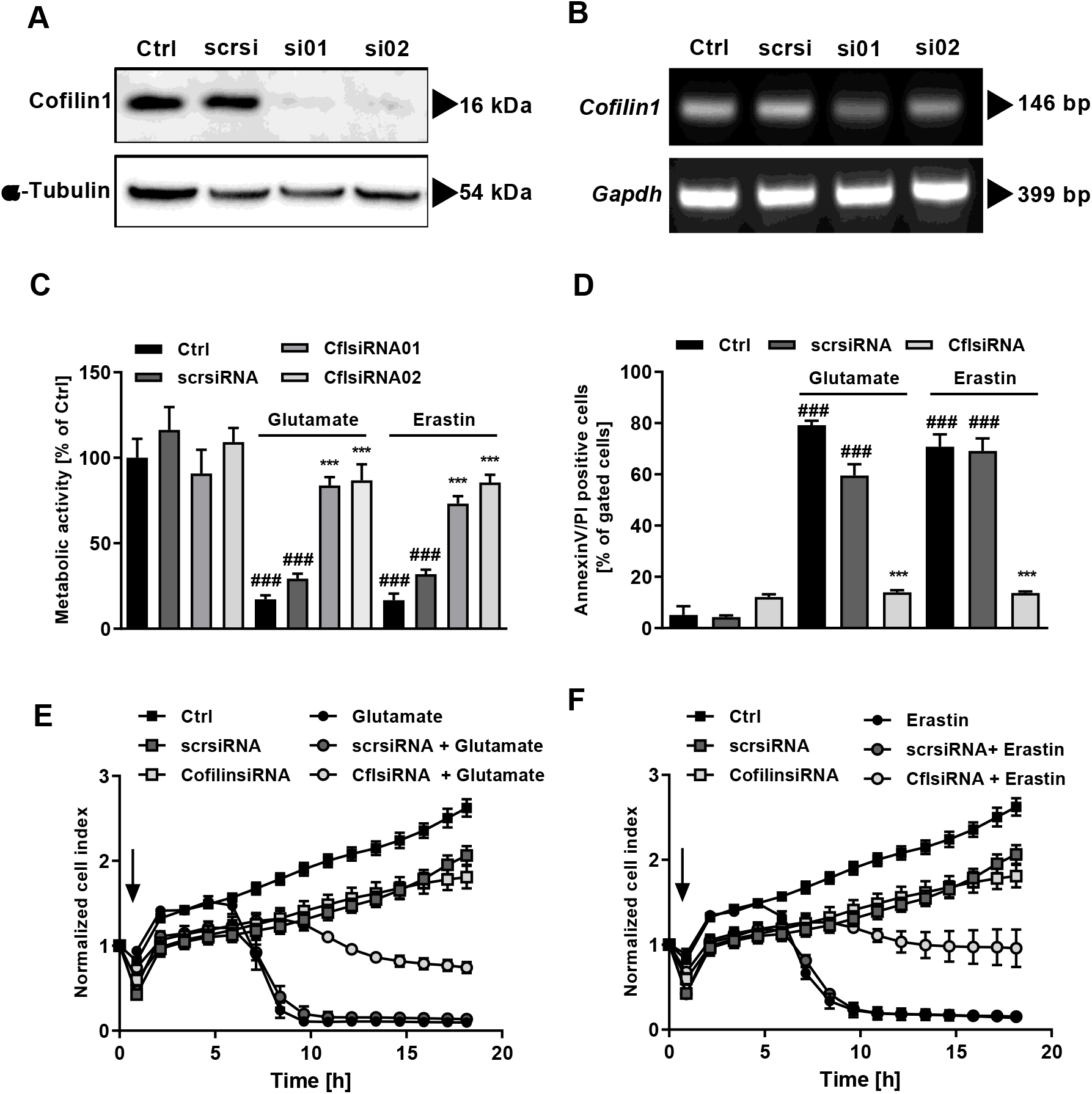
Cofilin1 depletion reveals sustained protection after glutamate or erastin exposure which occurs independent of lipid peroxidation and ROS accumulation. HT22 cells were transfected with two specific siRNAs against cofilin1 showing comparable effects. Unspecific scrambled siRNA (scrsi) was used as control. Transfection efficiency was analyzed after 48 h on **A** protein and **B** mRNA level. Protein levels were determined by Western blot using specific antibodies against cofilin1 and α-Tubulin as loading control. Transfection efficiency was also analyzed on *mRNA* level by RT-PCR. *Gapdh* was used as an internal control. **C** Cells were treated with 0.2 μM erastin or 2 mM glutamate for 16 h and were analyzed for proliferation/viability using MTT reagent. Both siRNA sequences conveyed comparable effects, therefore subsequent experiments were performed with siRNA1. Values are shown as mean + SD; n = 8 replicates. **D** AnnexinV and PI staining was conducted after 30 h of siRNA incubation and following 16 h of erastin (0.5 μM) or glutamate (2 mM) treatment. (mean + SD; 5,000 cells per replicate of n = 3 replicates). **E, F** xCELLigence measurement was performed after siRNA incubation for 30 h. The arrow indicates the time of erastin (0.75 μM) or glutamate (8 mM) application. Data are given as mean ± SD; n = 8 replicates. ### p<0.001 compared to untreated ctrl; ***p<0.001 compared to erastin- or glutamate-treated ctrl (ANOVA, Scheffé’s-test).

#### Cofilin1 acts downstream of lipid peroxidation and cellular reactive oxygen species formation, but upstream of mitochondrial demise

Oxidative distress is considered a major hallmark in oxytosis and ferroptosis [36, 37]. Peroxidation of lipids represents a hallmark in cell death cascades comprising reactive oxygen species downstream of glutathione depletion in models of oxytosis and ferroptosis [13, 15, 38]. To further validate the specific activity of cofilin1, lipid peroxidation was assessed using the fluorescent dye BODIPY C11 and flow cytometry; ROS were assessed using H_2_-DCF. Of note, both, erastin and glutamate treatment induced pronounced accumulation of lipid peroxides, which was not affected by siRNA-mediated cofilin1 knockdown (Fig. 3 A), indicating that lipid peroxidation occurs upstream of detrimental cofilin1 activation. Interestingly, cofilin1 depletion could not diminish the fluorescent DCF signal (Fig. 3 B), thus, in the applied model systems of oxidative cell death, cofilin1 activation occurs downstream of cellular ROS generation. Next, we used the MitoSOX marker that specifically detects mitochondrial superoxide. Mitochondrial involvement is widely considered as the ‘point of no return’ in cell death paradigms induced by millimolar doses of glutamate [10, 39, 40]. However, the relevance of mitochondrial damage in ferroptosis induction is controversially discussed [12, 13, 41, 42] and may depend on the experimental settings and, in particular, on the cell type. In this regard, it was essential to investigate possible links between ferroptotic signaling and mitochondrial demise. Intriguingly, both, glutamate and erastin challenge triggered massive mitochondrial superoxide accumulation, however, cofilin1 depletion completely blocked such mitochondrial superoxide formation (Fig. 3 C). To address the impact of cofilin1-knockdown on the mitochondrial membrane potential, TMRE staining and flow cytometry analysis were conducted. After glutamate or erastin exposure, the membrane potential was considerably impaired, and this was attenuated by cofilin1 knockdown (Fig. 3 D), thereby preserving mitochondrial integrity, represented by the mitochondrial membrane potential, which is crucial for proper energy storage upon oxidative phosphorylation processes [43]. Cofilin1 depletion without any further treatment neither impaired mitochondrial ROS generation, nor the mitochondrial membrane potential. Further, glutamate and erastin treatments resulted in an overall decline of ATP levels, measured by a luminescence-based approach. Depletion of cofilin1 was capable of partly rescuing the effect on ATP production indicating that, to a certain extent, the energy supply was preserved (Fig. 3 E). Massive mitochondrial calcium accumulation is considered as a detrimental prerequisite for mitochondrial impairment, eventually provoked by increased calcium-induced mitochondrial respiration, nitric oxide production and finally, loss of mitochondrial membrane integrity [44]. To address mitochondrial calcium alterations, Rhod-2 AM staining and flow cytometry measurements were performed. The results revealed, that cofilin1 depletion could significantly reduce the massive mitochondrial calcium accumulation following glutamate or erastin exposure (Fig. 3 F).

**Figure 3.**
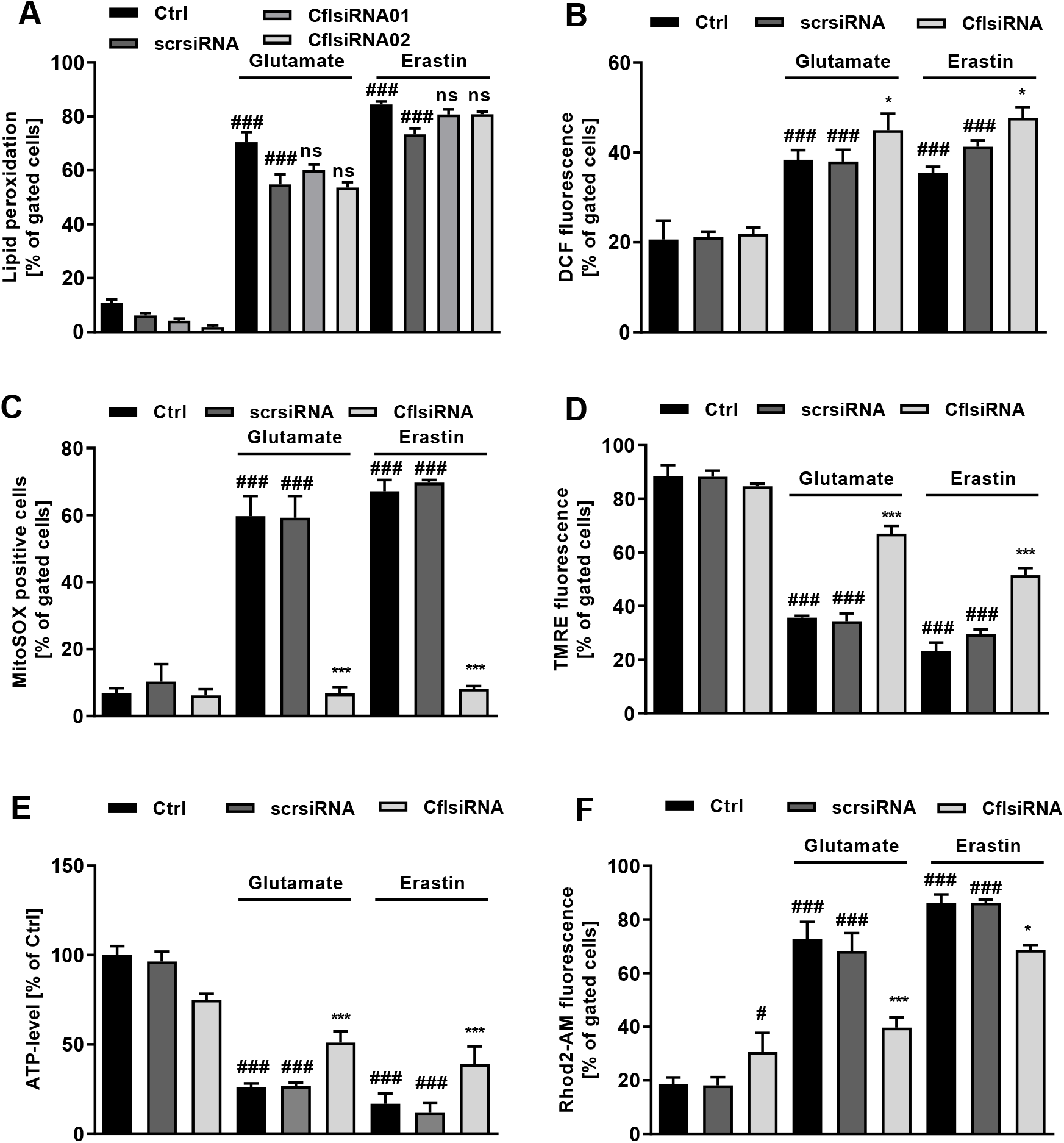
Cofilin1 silencing is effectively averting mitochondrial impairment in terms of glutamate or erastin toxicity. Prior to the described measurement, cofilin1siRNA was incubated for 30 hours. **A** Lipid peroxidation was determined 9 h after challenging the cells with 0.5 μM erastin or 5 mM glutamate with BODIPY fluorescent dye and subsequent FACS measurement. Data are given as mean + SD; 5,000 cells per replicate of n = 3 replicates. **B** The amount of ROS was measured after 0.8 μM erastin or 7 mM glutamate treatment for 10 h following H_2_DCF-DA staining and FACS analysis. Data are given as mean + SD; 5,000 cells per replicate of n = 3 replicates. **C** Mitochondrial superoxide accumulation was measured by MitoSOX staining and FACS analysis after 16 hours treatment with 0.5 μM erastin 4 mM glutamate. Data are presented as mean + SD; 5,000 cells per replicate of n = 3 replicates. **D** The mitochondrial membrane potential was evaluated by an appropriate cell permeant, positively-charged TMRE dye and following FACS analysis after 16 hours treatment with 1 μM erastin or 10 mM glutamate. Data are given as mean + SD; 5,000 cells per replicate of n = 3 replicates. **E** Cells were challenged for 8 hours with 0.7 μM erastin or 7 mM glutamate. ATP content was measured by luminescence-based measurement. Values are shown as mean + SD (n = 8 replicates. **F** Rhod-2 acetoxymethyl ester (Rhod-2 AM) was used to specifically measure mitochondrial calcium level after 16-hours treatment with 0.8 μM erastin or 8 mM glutamate. Values are projected as mean + SD; 5,000 cells per replicate of n = 3 replicates. Ctrl (control); scrsi (scrambled siRNA); Cfl1si (cofilin1 siRNA), # p<0.05 and ### p<0.001 compared to untreated ctrl, * p<0.05 compared to erastin- or glutamate-treated ctrl, ***p<0.001 compared to erastin- or glutamate-treated ctrl, ns = not significant (ANOVA, Scheffé’s-test).

With regard to energy metabolism, the Seahorse XFe96 Analyzer was used to determine both mitochondrial respiration by measuring the oxygen consumption rate (OCR) and glycolysis by measuring the extracellular acidification rate (ECAR). This measurement revealed complete inhibition of mitochondrial respiration and a diminished glycolysis rate after glutamate or erastin treatment in neural HT22 cells (Fig. 4 A-D). Cofilin1 downregulation partly preserved (Glutamate: *scrsi* 36.2±4.7 pmol/min → *Cfl1si* 53.6±8.1 pmol/min; Erastin: *scrsi* 35.7±7.7pmol/min → *Cfl1si* 50.0±5.5 pmol/min) mitochondrial respiration in both paradigms of cellular death (Fig. 4 A, C). In addition, the ability to generate energy by glycolysis was considerably maintained in cofilin1-knockdown cells in oxytosis and ferroptosis (Glutamate: *scrsi* 30.7±5.1 mpH/min → *Cfl1si* 69.9±2.8 mpH/min; Erastin: *scrsi* 28.2±6.3 mpH/min → *Cfl1si 68.6±2.8* mpH/min) (Fig. 4 B, D), suggesting a metabolic switch towards glycolytic-based energy production in cofilin1-deficient neurons exposed to oxidative stress. In this regard, correlation between OCR and ECAR illustrates the metabolic potential of the cells, measured under baseline and stressed conditions by FCCP injection (Fig. 4 E, F). Especially after glutamate or erastin treatment, metabolic bioenergetics underwent a mostly quiescent state in control conditions, whereas cells deficient of cofilin1 exhibited a considerably higher metabolic potential, indicating a functional energy production during oxidative stress (Fig. 4 E, F). Of note, under control conditions, cofilin1 knockdown itself significantly shifted the metabolic state (Fig. 4 E, F; Blue box).

**Figure 4.**
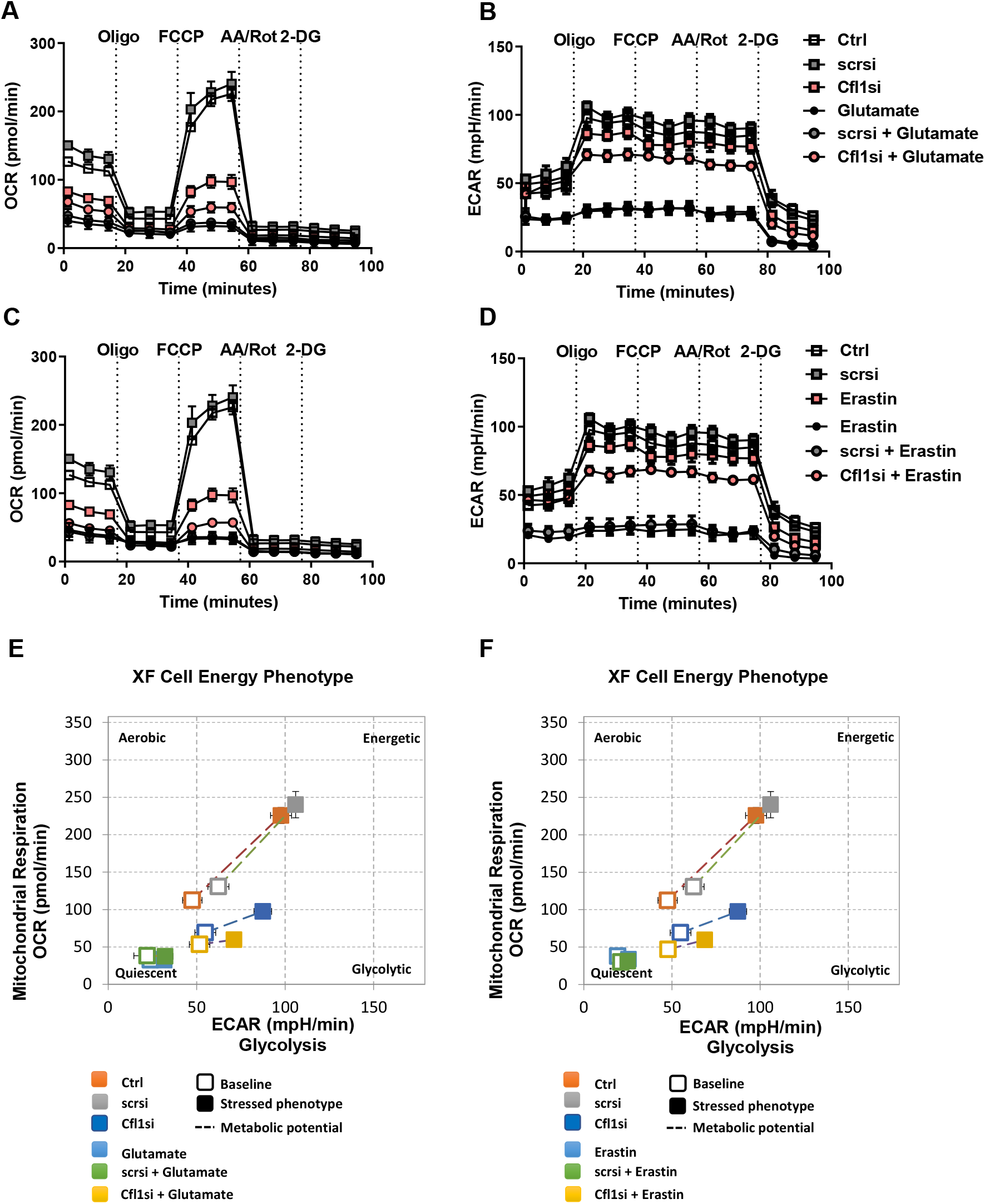
Cofilin1-deficient HT22 cells turn their energy production towards glycolysis after glutamate or erastin exposure. Cofilin1siRNA was transfected for 48 h. Afterwards, cells were damaged for 9 h with 0.5 μM erastin or 7 mM glutamate. **A, C** Afterwards, the oxygen consumption rate (OCR) and **B, D** the extracellular acidification rate (ECAR) were determined by a Seahorse XFe96 Analyzer. Data of 3 - 6 replicates per condition are shown as mean ± SD. Oligo (oligomycin); FCCP (carbonyl cyanide 4-(trifluoromethoxy)phenylhydrazone); AA (antimycin A) Rot (rotenone); 2-DG (2-deoxy-D-glucose). Ctrl (untransfected control); scrsi (scrambled siRNA); Cfl1si (cofilin1 siRNA). **E, F** The cell energy phenotype correlates the OCR and the ECAR of the cells at basal conditions (open dot) measured before the first compound was injected by the system and after FCCP injection, representing a stressed phenotype (filled dot). The displayed metabolic potential (dashed line) represents the capacity to meet the required energy demand under conditions of stress. Ctrl (control); scrsi (scrambled siRNA); Cfl1si (cofilin1 siRNA).

#### Primary cortical neurons deficient for cofilin1 showed enhanced resilience to glutamate excitotoxicity

To confirm the relevance of cofilin1 in neuronal cell death mechanisms, primary cortical neurons were isolated and used to analyze the impact of cofilin1 in the model of glutamate excitotoxicity. Therefore, primary cortical neurons of cofilin1^flx/flx^ mice referred to as control (Ctrl) cells and cofilin1^flx/flx, CaMKIIα-Cre^ regarded as cofilin1 knockout neurons, were used to evaluate the effect of cofilin1 in neuronal cell death mechanisms. Western blot analysis showed cofilin1 downregulation in Cre-expressing neurons after 30 days *in vitro* (DIV30) (Fig. 5 A). MTT assay was used to examine the remaining metabolic activity of cofilin1^-/-^ knockout cells after glutamate exposure in comparison to controls. In control cells, glutamate treatment led to a significant reduction of the metabolic activity, which was prevented by the NMDA-receptor antagonist MK801 (Fig. 5 B). In cofilin1-deficient neurons, the detrimental impact of glutamate was entirely abrogated, similar to the effects of MK801 (Fig. 5 B). Overall, excitotoxicity exerts detrimental effects on mitochondrial integrity and function, including deficits of ATP production as a consequence of impaired mitochondrial respiration. To gain further insight into the metabolic activity of primary neurons, we performed seahorse measurements. These assays revealed that the oxygen consumption rate, as an indicator of mitochondrial function, was decreased after glutamate exposure of Ctrl neurons under basal conditions and upon evoking maximal respiration by FCCP (Fig. 5 C, E, F). MK801 was able to prevent the detrimental impact of glutamate on basal and maximal OCR (Fig. 5 C, E, F). Similarly, mitochondria of cofilin1 knockout neurons were also significantly protected against glutamate-induced loss of basal and maximal respiration (Fig. 5 D, E, F), indicating that cofilin1 mediated mitochondrial damage in models of glutamate excitotoxicity in the cortical neurons.

**Figure 5.**
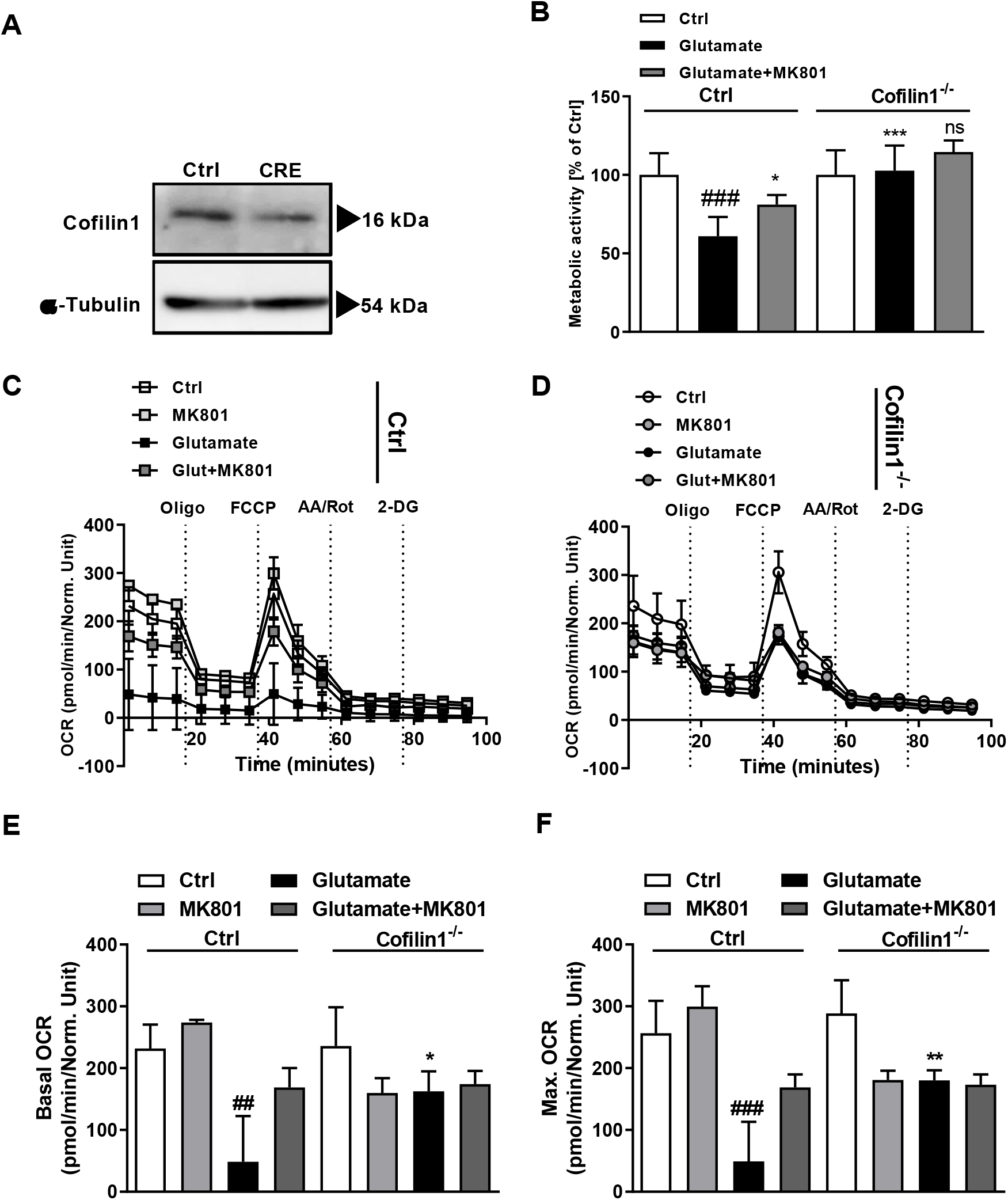
Cofilin1 knockout in primary cortical neurons reveals protection against glutamate-induced excitotoxicity. **A** Western Blot analysis was conducted after 30 *days in vitro* (DIV30) of control (Ctrl) neurons and cofilin1^flx/flx, CaMKIIα-Cre^ neurons (CRE). **B** Metabolic activity of DIV30 Ctrl and cofilin1^-/-^ neurons was determined by MTT assay after glutamate exposure for 24 h. MK801 cotreatment served as a control for NMDA-receptor blockade. Mean values + SD of n = 5 replicates are shown. **C** OCR of Ctrl and **D** cofilin1^-/-^ neurons was measured at 30 days in vitro after 25 μM glutamate challenge for 24 hours. 20 μM MK801 was applied simultaneously and served as a control for NMDA-R blockade. **E** Quantification of the basal OCR of Ctrl and cofilin1^-/-^ neurons, measured before the first compound was injected and **F** maximal OCR after FCCP injection of n = 3-5 replicates. Mean values ± SD are given. ns (not significant) compared to glutamate-treated cofilin1^-/-^ cells; ## p<0.01 and ### p<0.0001 compared to untreated ctrl, * p<0.05; ** p<0.01 and *** p<0.001 compared to glutamate-treated ctrl (ANOVA, Scheffé’s-test).

As mentioned before, the phosphorylation status of cofilin1 Ser3 is considered to be a decisive determinant for actin binding. Since the Rho-ROCK pathway was identified to activate LIM domain kinase 1 and 2 (LIMK1, 2) [45], a crucial cofilin1 negative regulator, the Rho activator CN03 was administered for induction of cofilin1 Ser3 phosphorylation and thus deactivation of the protein. To validate the effect of 1 μg/mL CN03 exposure on cofilin1 phosphorylation, Western blot was performed using a specific antibody against phosphorylated Ser3-cofilin1. As clearly demonstrated, cofilin1 was dephosphorylated after glutamate exposure, whereas CN03 preserved the phosphorylation status (Fig. 6 A). The effect of this manipulation was assessed in the MTT assay to quantify metabolically active cells. Interestingly, a 3-hour pretreatment with 1 μg/mL CN03 rescued the loss of metabolic activity induced by 24-hour exposure of glutamate (Fig. 6 B). This beneficial effect was comparable to the potent NMDA-receptor antagonist MK801 (Fig. 6 B).

**Figure 6.**
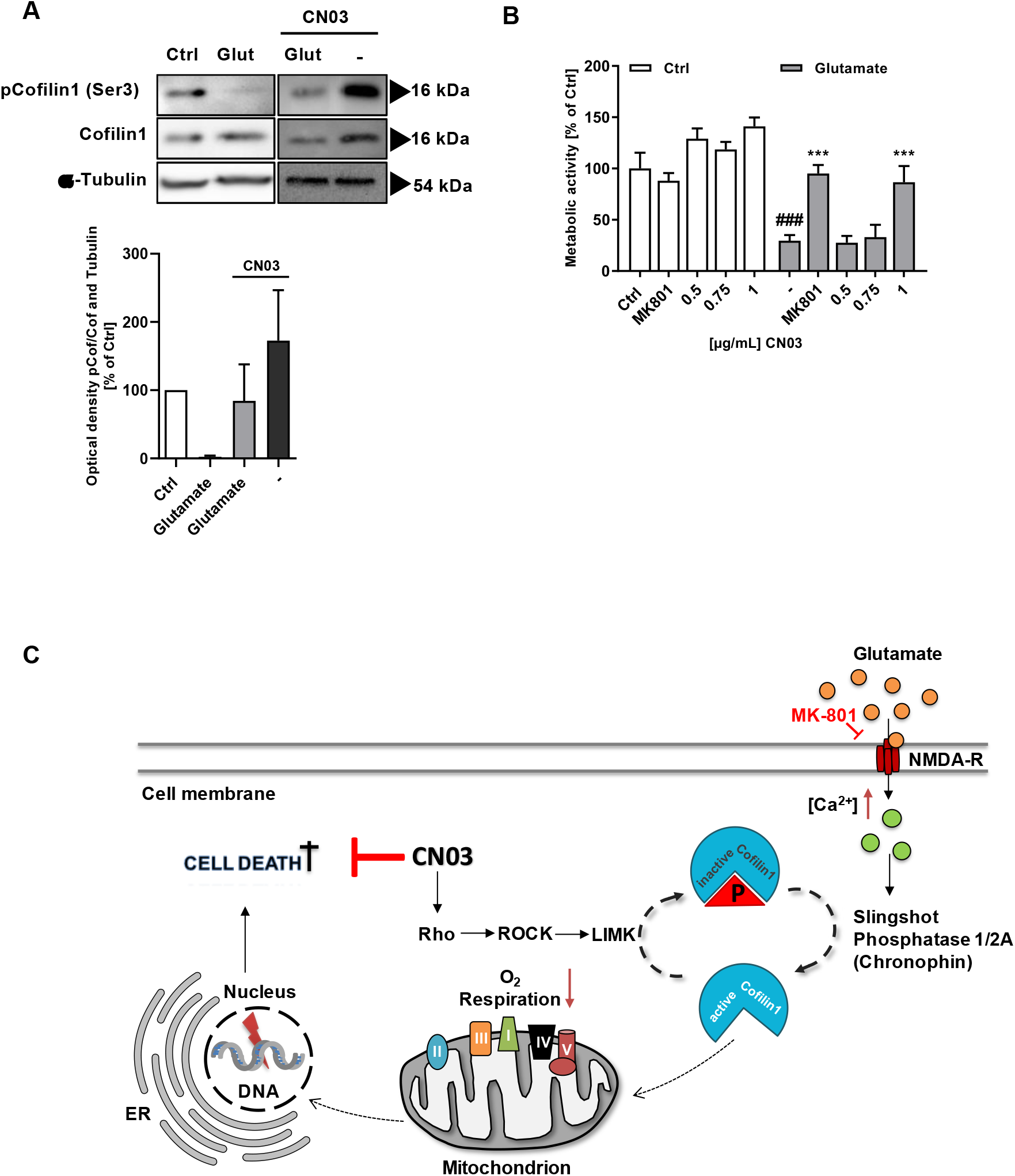
Enhancing cofilin1 phosphorylation enhances neuronal survival after glutamate exposure. **A** Western blot analysis of phosphorylated Ser3-cofilin1 was performed after 3 h pretreatment with 1 μg/mL CN03 and additional 24 h treatment with 25 μM glutamate. Quantification of the resulting signal was realized by densiometric analysis from n = 4 blots. The intensities of pCofilin1 (Ser 3) were compared to the cofilin1 signal and to α-Tubulin as a loading control and presented as mean + SD. Ctrl (control); Glut (glutamate). **B** Primary cortical neurons from wildtype E18 pubs were exposed to the indicated concentration of CN03 3 h prior to 25 μM glutamate treatment for 24 h at DIV9. Data from n = 6 are shown as mean + SD. ### p<0.001 compared to control; *** p<0.001 compared to glutamate-treated control (ANOVA, Scheffé’s-test). **C** Micromolar concentrations of glutamate stimulated excessive Ca^2+^ entry into neurons, a pathologic condition known as excitotoxicity. By application of CN03 protein, a known Rho-activator, cofilin1 is deactivated via ROCK-LIMK pathways thereby promoting neuronal protection by circumventing cofilin1 activation and mitochondrial demise. NMDA-R (N-methyl-D-aspartate receptor); ER (endoplasmic reticulum); [Ca^2+^] (intracellular calcium concentration); P (Ser 3-phosphorylation); ROCK (Rho-associated serine/threonine kinase); LIMK (LIM kinase); DNA (deoxyribonucleic acid); roman numerals representing complex I-V of the respiratory chain.

#### Administration of recombinant cofilin1 on isolated mitochondria impaired the mitochondrial membrane potential and respiration

Besides the well-established function on F-actin dynamics, cofilin1 has been linked to oxidative cell death, e.g. induced by the oxidant taurine chloramine [22] or H_2_O_2_ [46, 47]. Cofilin1 possesses several cysteine residues essential for the quaternary structure of the protein by forming intra- or intermolecular disulfide bonds. In human cofilin1, four crucial cysteines have been described prone to oxidation at positions 39, 80, 139 and 147 [24]. Dephosphorylation of Ser3 and oxidation of the aforementioned cysteine residues are considered as crucial prerequisites for mitochondrial localization after apoptosis induction [22]. To specifically address the impact of WT cofilin and specific mutants (2Cys: Cys139Ser/Cys147Ser; 4Cys: Cys39Ser, Cys80Ser, Cys139Ser, Cys147Ser), we cloned different constructs, expressed the proteins in E. coli and purified it using the IMAC principle. Proteins were re-buffered into PBS and the impact of Cys mutations on the thermal stability was analyzed at 600 nm using a UV-spectrophotometer. The 2CysSer mutant showed a similar stability as the WT protein (Supplementary Fig. 1). The 4CysSer mutant was less stable than the other proteins. However, all recombinant proteins were stable at the temperature of 37 °C that was used for all experiments on isolated mitochondria (Supplementary Fig. 1). The impact of recombinant cofilin1 was analyzed on mitochondrial superoxide formation, mitochondrial membrane potential and mitochondrial respiration. Mitochondria isolated from mouse brain were incubated with or without the respective cofilin1 variants under basal, oxidized (H_2_O_2_) or reduced conditions (DTT). Strikingly, the reduced form of WT cofilin1 had no impact on the membrane potential, whereas application of WT cofilin1 either in the natural form or in the oxidized state decreased the mitochondrial membrane potential. Further, the Cys139/147Ser mutation as well as conversion of all four cysteines to serine completely abolished the effect of cofilin1 on isolated mitochondria (Fig. 7 A). The chemical ionophore CCCP was used to demonstrate the maximal effects of mitochondrial membrane collapse measured by TMRE staining (Fig. 7 A).

**Figure 7.**
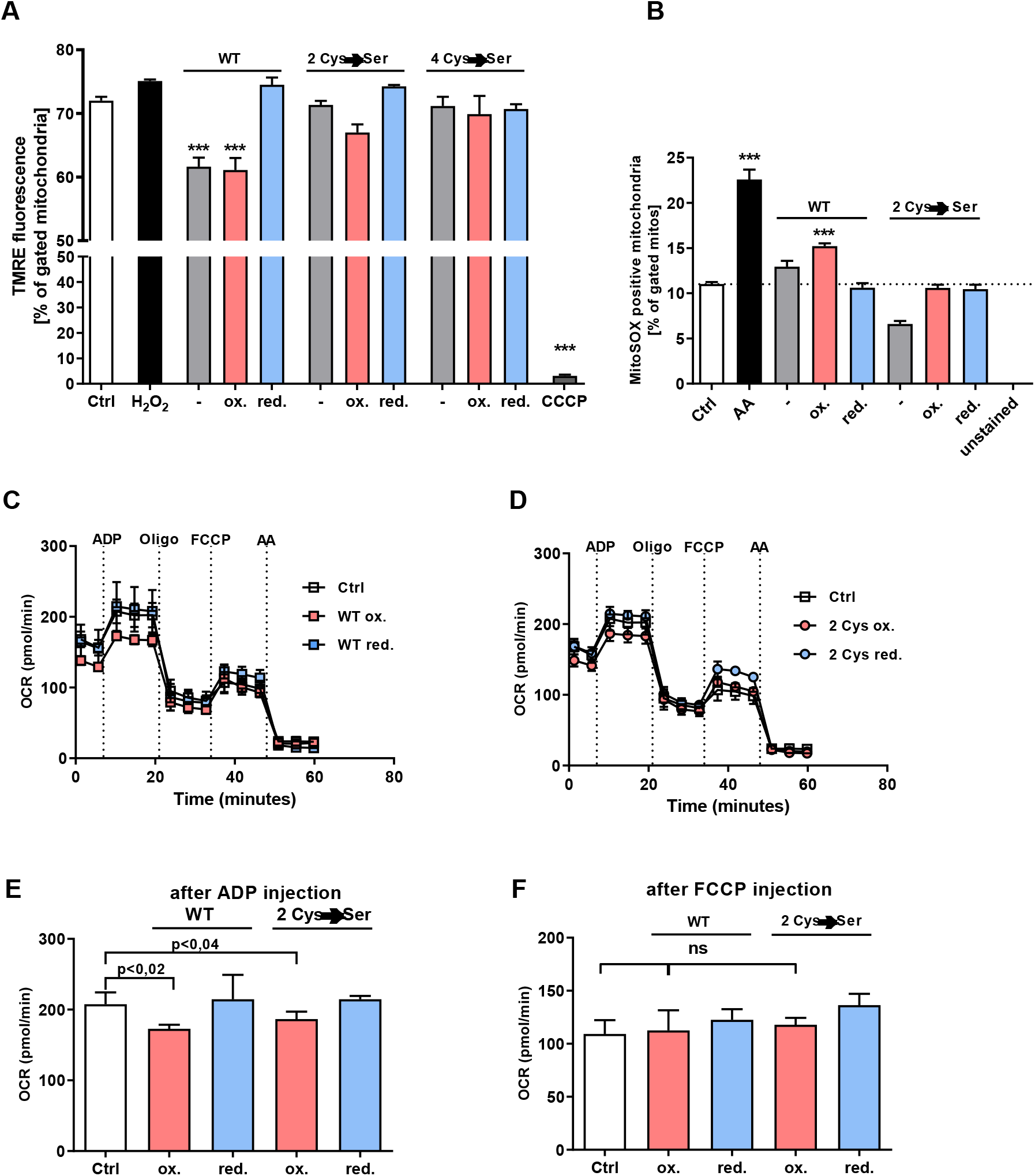
Direct effects of recombinant, oxidized cofilin1 impair the mitochondrial membrane potential and respiration. **A** Recombinant cofilin1 protein was applied in the native, the oxidized (100 μM H_2_O_2_) or in the reduced form (10 mM DTT). 75 μg mitochondria were incubated with 10 μg protein for 20 minutes at 37 °C and finally stained with TMRE (1:1,000). 50 μM CCCP served as a positive control. 10,000 total events were measured and shown as mean + SD (n = 3 replicates). WT (wildtype cofilin1 protein); 2Cys (Cys139/Cys147 → serine mutation); 4Cys (39, 80, 139, 147 → serine mutation) *** p<0.001 compared to ctrl (ANOVA, Scheffé’s-test). **B** 75 μg mitochondria were incubated with 10 μg protein for 20 minutes at 37 °C and finally stained with MitoSOX red fluorescent dye (1:1,000). 10 μM Antimycin A (AA) served as a positive control. 10,000 total events were measured and shown as mean + SD (n = 3 replicates). WT (wildtype cofilin1 protein); 2Cys (Cys139/Cys147 → serine mutation); ***p<0.001 compared to ctrl (ANOVA, Scheffé’s-test). **C** 10 μg mitochondria per well were incubated with the WT protein or **D** with the 2 Cys mutant either in the native form, the oxidized form (100 μM H_2_O_2_) or in the reduced form (10 mM DTT) for 30 minutes at 37 °C and were administered to the Seahorse Analyzer. Mean + SD (n = 5-9 replicates). **E** Quantification of mitochondrial activity was conducted with the values delivered after the injection of ADP as a substrate for the OXPHOS phosphorylating capacity. **F** FCCP uncouples the oxygen consumption from ATP production and is used to assess maximal respiratory activity. ns (not significant); WT (wildtype cofilin1 protein); 2 Cys (Cys139/Cys147 → serine mutation); (p values were calculated by ANOVA, Scheffé’s-test).

To further elucidate the impact of the WT protein and the 2Cys-cofilin1 mutant on isolated mitochondria, mitochondrial superoxide generation was measured using MitoSOX staining. The maximal effect of mitochondrial superoxide generation was evoked by antimycin A treatment, a potent complex III-inhibitor of the respiratory chain (Fig. 7 B). The oxidized form of the WT protein also induced a significant burst of mitochondrial superoxides, whereas the reduced WT protein and the 2Cys-mutant generated comparable ROS levels to mitochondria of the untreated control group (Fig. 7 B). The effect of the serine mutants on mitochondrial integrity and superoxide generation was therefore mainly attributed to the cysteine residues at position 139 and 147.

Finally, to understand the effect of recombinant cofilin1 at a functional level, mitochondrial bioenergetics were evaluated using the Seahorse XFe Analyzer. The mitochondrial assay buffer (MAS) contained succinate to specifically assess complex II-driven respiration and rotenone to prevent reverse electron flow. ADP injection allows for specific calculation of the phosphorylating capacity to produce ATP. The analysis of the ADP-driven mitochondrial activity revealed a significant impairment (p<0.02) of the measured OCR of mitochondria challenged with the oxidized WT cofilin1 protein compared to the control condition (Fig. 7 C, D, E). The oxidized form of the 2Cys mutant had a slightly minor derogating impact on the ADP-dependent respiration compared to the WT cofilin1 protein (WT p<0.02 vs. 2Cys p<0.04), which was completely reversed by reduction of these cysteine residues (Fig. 7 D, E, F). Oligomycin, a potent ATP synthase inhibitor (complex V), allowed for estimating the proton leak across the inner mitochondrial membrane, which was not apparently affected after application of either the oxidized or reduced form of the recombinant protein (Fig. 7 C, D). Injection of the uncoupler FCCP disrupts the proton gradient to facilitate maximal respiration. Again, this state of mitochondrial respiration was not compromised by either the WT cofilin1 or the 2Cys cofilin1 mutant (Fig. 7 F).

### Discussion

The present study identified a pivotal role of cofilin1 upstream of mitochondrial damage in oxidative cell death induced by glutamate or erastin and in models of glutamate excitotoxicity in primary cortical neurons. Here, we demonstrate that cofilin1 downregulation in neuronal HT22 cells by specific cofilin1-targeting siRNA or by genetic deletion in primary neurons, exerts protective effects on mitochondrial function and cellular resilience. We have shown that cofilin1 deficiency affects mitochondrial superoxide accumulation, mitochondrial membrane potential, mitochondrial calcium accumulation, mitochondrial respiration, and ATP generation.

Mitochondrial superoxide production by complex I and III occurs during respiration. Moreover, it can either be a result of complex I defects, genetic abnormalities or excessive mitochondrial calcium admission [48–50]. Massive mitochondrial calcium gathering induces the opening of the mitochondrial permeability transition pore and subsequent loss of the mitochondrial membrane potential leading to cell death (reviewed in [51]). Our results emphasize that cofilin1 depletion can reduce massive mitochondrial calcium overload which may account for the protective mechanism upon cofilin1 depletion. In this regard, possible indirect effects by alteration of the actin cytoskeleton might be conceivable to impact mitochondrial calcium uptake, as previously discussed for INF2-knockout cells [52]. Mitochondrial ROS is especially detrimental for mitochondrial function, as it leads to mitochondrial DNA damage and subsequent impaired oxidative phosphorylation (OXPHOS) [53–55]. Therefore, preventing mitochondrial ROS accumulation by cofilin1 depletion is an efficient intervention point to retrieve neurons under pathophysiological conditions, as imposed here through glutamate- or erastin treatment.

The precise point of the detrimental action of cofilin1 was ascertained upstream of mitochondrial demise, but downstream of lipid peroxidation. Further, AnnexinV/PI staining, xCELLigence measurement and MTT-assay revealed that cell death was also attenuated by interfering with cofilin1 expression. Mitochondria are regarded as the point of no return upon cell death induction [56]; therefore, cofilin1 inhibition is efficient in preventing cell death in these paradigms of oxidative damage since cofilin1 is an important key player in death signaling upstream of mitochondria.

The conditional mouse model of genetic cofilin1 deletion in neurons is a powerful strategy to study cofilin1 involvement under conditions of glutamate excitotoxicity in primary neurons. Excessive glutamate accumulation is a hallmark of several neurodegenerative diseases, stroke and brain trauma [57, 58]. Many hallmarks of neuronal loss are already identified to facilitate the development of possible treatment strategies for these diseases. However, how different cell death mechanisms are involved and at which point they converge is not completely understood. Cofilin1 was identified as a key player in different neurological diseases, e.g. in Alzheimer’s or Parkinson’s disease [59–62]. Recent findings also imply a role for cofilin1 in ischemic brain damage [63, 64]. In this paradigm, cofilin1 phosphorylation was able to prevent detrimental cofilin-actin rod formation, thereby improving mitochondrial transport failure induced by oxygen and glucose deprivation [63]. In our study, CN03-induced phosphorylation also impressively protected neurons from glutamate-induced cell death revealing a potential target for future treatment strategies, conceivable for a variety of neurological disorders, such as stroke [63] or autism-like deficits [65]. A putative mechanism involves Rho activation by CN03, thereby phosphorylating cofilin1 via ROCK-LIMK pathways and finally promoting neuronal protection by circumventing cofilin1 activation and mitochondrial demise (Fig. 6 C).

Underlining the strong effect of our loss-of-function-approach, evidence exists, showing a direct effect of cofilin1 by binding to mitochondria [22, 66, 67]. In this regard, it was demonstrated that cofilin1 gains activity to translocate to mitochondria upon oxidation by taurine chloramine, a physiological oxidant derived from neutrophils [22, 34]. In line with these results, our data suggest that cofilin1 is an important mediator upon oxidative distress, as cofilin1 depletion was sufficient to block mitochondrial damage and, thereby, the oxidative cell death cascades. In order to distinguish potential indirect actin-based cofilin1 effects from those which are directly mediated by cofilin1, isolated mitochondria were incubated with recombinant cofilin1 protein. The functional measurements revealed a direct detrimental effect of cofilin1 on mitochondrial integrity and respiration. First hints of direct cofilin1-mediated impacts on mitochondria were given by Chua and coworkers in 2003 [23]. A few years later, four cysteine (39/80/139/147) and three methionine residues were identified, which are prone to oxidation, but only the oxidized cysteines were linked to mitochondrial demise [22]. In particular, a detrimental role for the oxidized form of cofilin1 upstream of mitochondria was unraveled in cell death models induced by the oxidants H_2_O_2_ or taurine chloramine (TnCl) [22, 68]. Under these conditions, cofilin1 can translocate to mitochondria and induce mitochondrial swelling, cytochrome c release and open the mitochondrial permeability transition pore (mPTP). This mitochondrial transactivation of the protein was even observed under basal conditions without any further stimulus when the cells express the oxidation-mimetic glycine residues at position 39 or 80, respectively [68]. Apparently, cysteines do not only serve as redox sensors and mediate redox signaling, they are also crucial for the correct structural protein formation and interaction. Especially Cys39 and Cys80 were described to form intramolecular disulfide bonds and their oxidation eventually led to protein dephosphorylation (Ser3) after oxidation due to sterical effects [24]. Cys139 and Cys147 are able to form both, intra- and intermolecular disulfide bonds, thus presenting a prerequisite for cofilin1 oligomerization [69]. In the present study, specific mutations of either two (Cys139/147) or all four cysteine residues of the recombinant protein were realized to address the question which specific cysteine residues contribute to the deleterious effects of the protein after oxidation. Specific evaluation of mitochondrial parameters after incubation of isolated mitochondria with cofilin1 facilitated insights into the direct mechanism of cofilin1 without any other cellular components. Intriguingly, the wildtype form of cofilin1 significantly decreased the mitochondrial membrane potential in isolated mitochondria. This detrimental effect on mitochondrial membrane integrity was attenuated when cofilin1 residues at position 139 and 147 were mutated to the non-oxidizable amino acid serine; and this effect was completely averted when all four cysteine residues were substituted by serine. These data suggest that oxidation of cofilin1 lead to a significant impairment of mitochondrial integrity and function. Accordingly, mitochondrial ROS accumulation was enhanced by the oxidized form of cofilin1 and, in line with the TMRE measurements, the 2Cys mutant did not exert mitochondrial ROS formation. These findings are in line with findings by Klamt *et. al* who evaluated the Cys139/147 mutant in a cellular environment by transfection of respective cofilin1-mutated plasmids [22].

Further, evaluation of the mitochondrial respiration revealed, that the wildtype form of cofilin1 impaired the oxygen consumption upon ADP injection, an indicator of complex II, III and V-driven respiration. Although the mutation of Cys139 and 147 still led to a decrease of mitochondrial respiration, the effect was less pronounced compared to wildtype cofilin1.

In conclusion, the deleterious effect of cofilin1 was attenuated if either all cysteine residues of the protein were substituted by the non-oxidizable serine, or if cysteine residues at position 139 and 147 were mutated, indicating that both positions, Cys139 and Cys147, are crucial in mediating the direct damaging impact on mitochondria. However, the effects of the oxidized form of cofilin1 were always less pronounced compared to the positive controls (Antimycin A for mitochondrial-derived ROS and CCCP as an uncoupler to induce collapse of the mitochondrial membrane potential). A possible explanation is delivered by Liu and coworkers who demonstrated that cofilin1 needs the interaction with p53 for strong impacts on mitochondrial function [67]. Finally, our data demonstrate a striking effect of cofilin1 in cell death mechanisms linked to oxidative distress, underlining that cofilin1 acts as a redox sensor in cell death mechanisms comprising oxytosis and ferroptosis. However, it is tempting to speculate that thiol switches of cofilin1 generally have also regulatory functions in physiological signal transduction pathways. Thus, our data suggests, that interfering with cofilin1’s activity by pharmacological inhibition or imposing cofilin1 phosphorylation at serine residue 3 could provide new potential therapeutic strategies for neurodegenerative diseases in the future.

## Acknowledgements

L.H., C.C. and M.R. were supported by the DFG Research Training Group 2213 ‘Membrane Plasticity in Tissue Development and Remodeling’. This work was further supported by Mitonetwork funded through the Flexifunds by the Forschungscampus Mittelhessen FCMH and ACAciA consortium. We thank Dr. Walter Witke (University of Bonn) for providing floxed cofilin1 mice.

---

**Figure 1 S.**
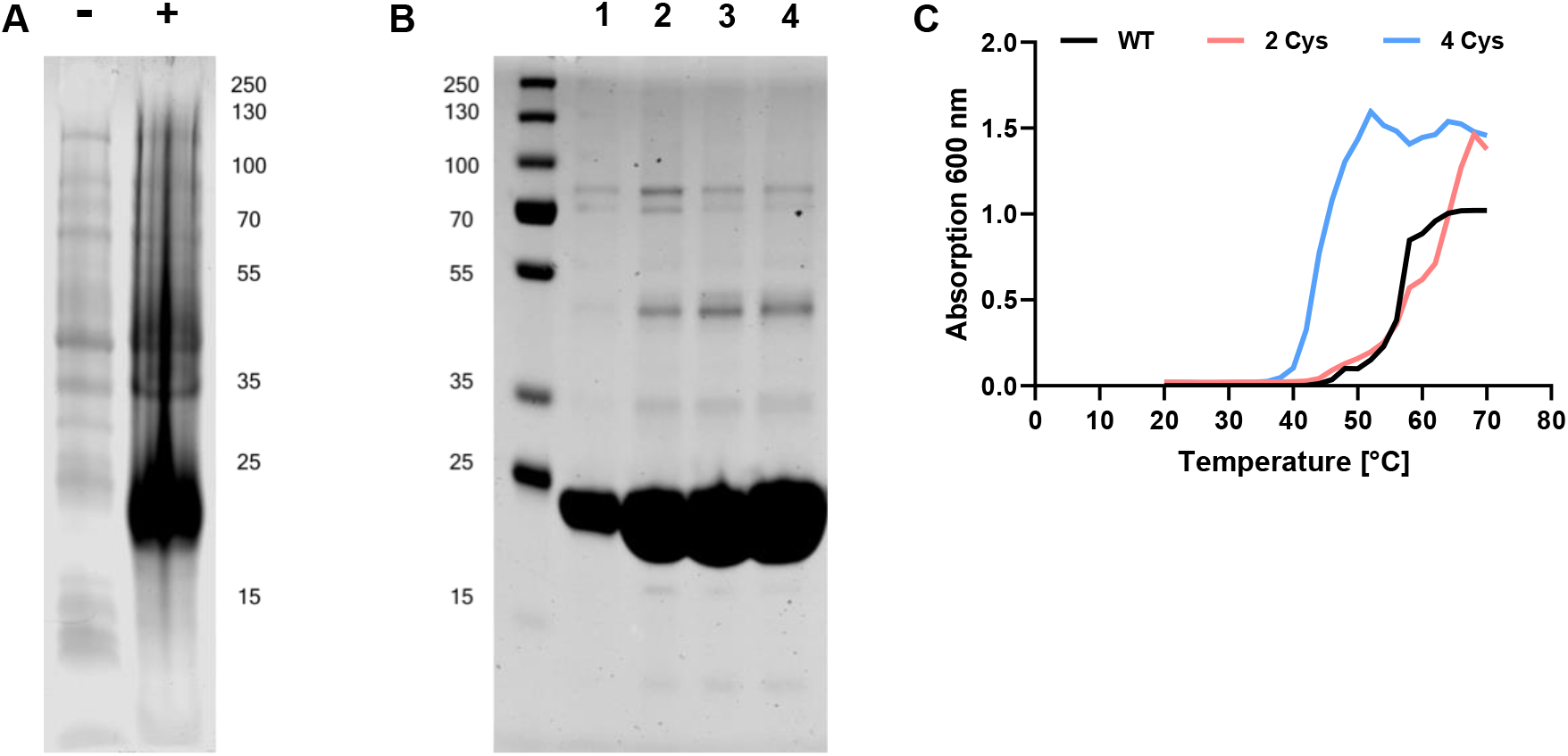
Expression, purification and thermal stability of recombinant cofilin1. Recombinant WT, 2Cys→ Ser and 4Cys→ Ser Cofilin-1 were expressed in E. coli and purified using the IMAC principle. **A** Expression samples prior (-) and after induction (+) with Isopropyl β-D-1-thiogalactopyranoside (IPTG). **B** Purified proteins were analyzed by SDS-Page and Coomassie staining. The exemplary illustration shows WT Cofilin1 elution fractions 1-4 with a molecular weight of 18 kDa. **C** Proteins were incubated at increasing temperatures and thermal stability was determined photometrically at 600 nm.

